# Comparative Analysis of Sorghum EMS Mutants and Natural Populations

**DOI:** 10.1101/2021.06.06.447271

**Authors:** Liya Wang, Anna Lipzen, Zhenyuan Lu, Junping Chen, Xiaofei Wang, Marcela K. Tello-Ruiz, Kerrie Barry, Jenny Mortimer, Doreen Ware, Zhanguo Xin

## Abstract

To build a large-scale genomic resource for functional validation of sorghum genes through EMS-mutagenized BTx623 seeds, we deep sequenced (30-60X) an additional 445 phenotyped EMS mutant lines. 4.2 million EMS mutations are called with nearly 36,800 mutations that could have a disruptive effect on functions of over 15,500 genes. Combining variants carried by both the natural population and previous EMS efforts, over 69% of sorghum coding genes (23644) are now presented with one or more mutations that are, or are predicted to be, disruptive to their functions. Our results show that the EMS population carries more significant mutations but less in each sample than the natural population, which makes it more powerful in elucidating sorghum gene functions on a large scale and requiring less work in validation of candidate causal genes. We have made the data available through two ways, one is the integration with the BSAseq workflow that supports retrieving independent EMS samples carrying the same genes with significant mutation for complementary testing, and the other is a web application for directly querying genes with significant mutations on SciApps.org.

## Introduction

Compared with traditional map-based cloning, molecular identification of allelic variation responsible for the phenotypes of plants with induced mutants, known as forward genetics, has been much more effective in demonstrating gene functions. For introducing point mutations, ethyl methanesulfonate (EMS) mutagenesis has been widely used for discovering gene functions for Arabidopsis (Page and Grossniklaus 2002), rice (Abe et al. 2012), and sorghum [EMS1 (Addo-Quaye et al. 2017; Addo-Quaye et al. 2018), and our previous effort, EMS2 (Jiao et al. 2016)]. In this study, we deep-sequenced an additional 445 EMS pools to further boost the coverage of genes with significant mutations. We also compared the statistics of EMS data with recently released large natural population data (Lozano et al. 2021). Genes with significant mutations from both populations were integrated and are available through the SciApps platform.

## Materials and Methods

### EMS treatment and DNA sample preparation

EMS treatment was performed as described before (Xin et al. 2008).

### Variation analysis

Whole-genome sequencing of the 445 EMS-treated lines was performed on Illumina HiSeq 2000 sequencing system. The sequencing depth of each line varies with the majority of them falling between 30X - 60X (Figure S1). All reads were aligned using the Burrows-Wheeler Alignment (BWA) tool (Li and Durbin 2009) to the sorghum reference genome v3.1 available in Phytozome (McCormick et al. 2018). Variant calling was performed with Samtools and Bcftools (Li and Durbin 2009; Li et al. 2009). Variants overlapping with the ms8 founder line (Wang, Lu, Regulski, et al. 2020), the parental BTx623 line, were removed to eliminate background mutations. High-confidence EMS SNPs were selected using the following criteria: (1) the nucleotide change is GC → AT; (2) the SNP is supported by at least 5 reads; (3) the mutation has an allele frequency <= 5%.

1 out of the 445 lines (ARS1162) is removed since it carries at least 2.5 times more EMS SNPs than the majority of other lines (54K vs 5K-20K), which is an indication of contamination. For the remaining 444 lines, cross contaminations were checked via the overlapping of EMS SNPs between any pair of lines (Figure S2). Results show that over 99.99% of the pairs have < 5% overlapping EMS mutations and over 97% have <1% identical EMS mutations shared between any two lines. However, we do see very rare highly overlapping EMS SNPs between samples with adjacent ids (along the diagonal line), which might be an indication of biological contamination. We decided to keep both samples in such a situation since they both may contain the significant mutations and the true causal genes could still be easily verified.

Functional annotation of SNPs was performed using SnpEff (Cingolani et al. 2012) and deleterious mutations were predicted using SIFT 4G (Vaser et al. 2016). Both were done on the SciApps platform (Wang et al. 2018; Wang, Lu, delaBastide, et al. 2020).

### Data availability

EMS variant data are available from the SorghumBase’s CyVerse repository: https://datacommons.cyverse.org/browse/iplant/home/shared/SorghumBase/ems. We built a Shiny app for checking the presence of a single gene with any significant mutations: https://data.sciapps.org/shiny/ems/?gene_id=Sobic.002G221000. The app is built into the BSAseq workflow (Wang, Lu, Regulski, et al. 2020) for facilitating complementary testing of true causal genes. A web-based application is also provided for checking a list of genes against both the EMS population and the natural population: https://www.sciapps.org/app_id/queryEMS-0.0.1.

## Results

### Population sequencing and variant analysis

Because many M2 plants are sterile, mutagenized M1 seeds were propagated to M3 seeds through single-seed descent, and genomic DNA was prepared from leaf samples pooled from 20 individual M3 plants. A total of 445 M3 families were randomly selected for whole-genome sequencing. To capture the majority of the heterozygous EMS mutations, we chose to deep sequence each family (30X-60X). Although the number of EMS SNPs does not show strong correlation with the sequencing depth (Figure S1), with deep sequencing we had a higher chance to capture the majority of the heterozygous SNPs presented in the M3 families and more confidence in the called variants.

### Functional annotation of the EMS-induced mutations

We used SnpEff (Cingolani et al. 2012) to annotate the SNPs generated by EMS treatment or present in the natural population. The effect of nonsynonymous substitutions were predicted by SIFT 4G (Vaser et al. 2016). To reduce the number of false-positive predictions made by SIFT, we also used the genomic evolutionary rate profiling (Davydov et al. 2010) or GERP score, which identified 9.49% (64.9 Mb) of the sorghum genome (Valluru et al. 2019) as evolutionarily constrained (GERP > 0).

In this study, we use four categories to classify a significant mutation: (1) predicted, with SIFT score < 0.05, median_info < 3.25, and GERP score >= 2. A median_info value greater than 3.25 implies that there are not enough sequences to make the SIFT score prediction; (2) knockout, include high impact mutations that are annotated by SnpEff as SPLICE_SITE_ACCEPTOR, SPLICE_SITE_DONOR, STOP_GAINED, or START_LOST; (3) stop-gained, constituting the majority of the knockout mutations; (4) significant, the union of all three categories above. A gene is only counted once in Table 1 if it has more than one significant mutation. In total, over 69% of the protein-coding genes are now presented in one or more of the 1681 lines carrying a significant mutation.

**Table 1.** Summary of SNPs and impacts on genes for EMS mutants and natural populations. The numbers in the parenthesis are the average numbers per sample (for SNPs) and percentage of total 34,118 sorghum v3 protein coding genes (for genes)

|  | **EMS1** | **EMS2** | **EMS3** | **All EMS** | **NP** | **All** |
| --- | --- | --- | --- | --- | --- | --- |
| Ref. | Addo-Quaye, 2018 | Jiao, 2016 | - | - | Lozano, 2021 | - |
| # samples | 486 | 252 | 444 | 1182 | 499 | 1681 |
| # SNPs | 2.55M<br>(5247) | 1.74M<br>(6905) | 4.17M<br>(9392) | 8.41M<br>(7115) | 13.1M<br>(26253) | 21.4M<br>(12731) |
| # predicted significant SNPs | 19563<br>(40.2) | 15870<br>(63.0) | 26127<br>(58.8) | 60652<br>(51.3) | 13872<br>(27.8) | 74288<br>(44.2) |
| # knockout SNPs | 7621<br>(15.7) | 6180<br>(24.5) | 10667<br>(24.0) | 24207<br>(20.5) | 7465<br>(15.0) | 31551<br>(18.8) |
| # stop-gained SNPs | 5334<br>(11.0) | 4505<br>(17.9) | 7961<br>(17.9) | 17606<br>(14.9) | 4613<br>(9.2) | 22153<br>(13.2) |
| # significant SNPs | 27175<br>(55.9) | 22039<br>(87.5) | 36785<br>(82.8) | 84831<br>(71.8) | 21333<br>(42.8) | 105800<br>(62.9) |
| #predicted significant genes | 9595<br>(28.1%) | 8084<br>(23.7%) | 10294<br>(30.2%) | 13444<br>(39.4%) | 6998<br>(20.5%) | 13808<br>(40.5%) |
| # knockout genes | 6431<br>(18.9%) | 5187<br>(15.2%) | 8428<br>(24.7%) | 15033<br>(44.1%) | 5497<br>(16.1%) | 17663<br>(51.8%) |
| # stop-gained genes | 4689<br>(13.7%) | 3907<br>(11.5%) | 6610<br>(19.4%) | 12077<br>(35.4%) | 3550<br>(10.4%) | 14073<br>(41.2%) |
| # significant genes | 13642<br>(40.0%) | 11372<br>(33.3%) | 15532<br>(45.5%) | 21384<br>(62.7%) | 11557<br>(33.9%) | 23644<br>(69.3%) |

Table 1 shows that the number of significant SNPs per line is similar between EMS2 and EMS3, and both are higher than EMS1 in each category. The difference might be due to the fact that the EMS1 lines are individual M3 plants, not a mixture of 20 plants as in EMS2 and EMS3. All EMS lines have a higher number of significant SNPs than the natural population, with EMS2 and EMS3 both having two times more stop-gained or total significant mutations per line than the natural population, though the total EMS mutations (8.41M) is less than the total natural mutations (13.1M).

### Statistics of genes with significant mutations

Table 1 shows that EMS3 has the highest coverage of genes with significant mutations in each category than EMS1, EMS2, and the natural population (herein referred to as NP). Specifically, as shown in Figure 1, EMS3 adds 4902 new genes with knockout mutations, in which there are 4335 new genes with stop-gained mutations. The number of new genes from EMS3 with mutations predicted to be significant (1601) or total significant mutations (3943) are smaller since we are only counting a gene once when there are multiple mutations present in the same gene and there is a much larger overlap among EMS datasets in these two categories (Figure 1A and 1C) than knockout or stop-gained mutations (Figure 1B and 1D). This is also true when compared with the natural population (Figure S3), where the natural population also adds new genes with significant mutations.

**Figure 1.**
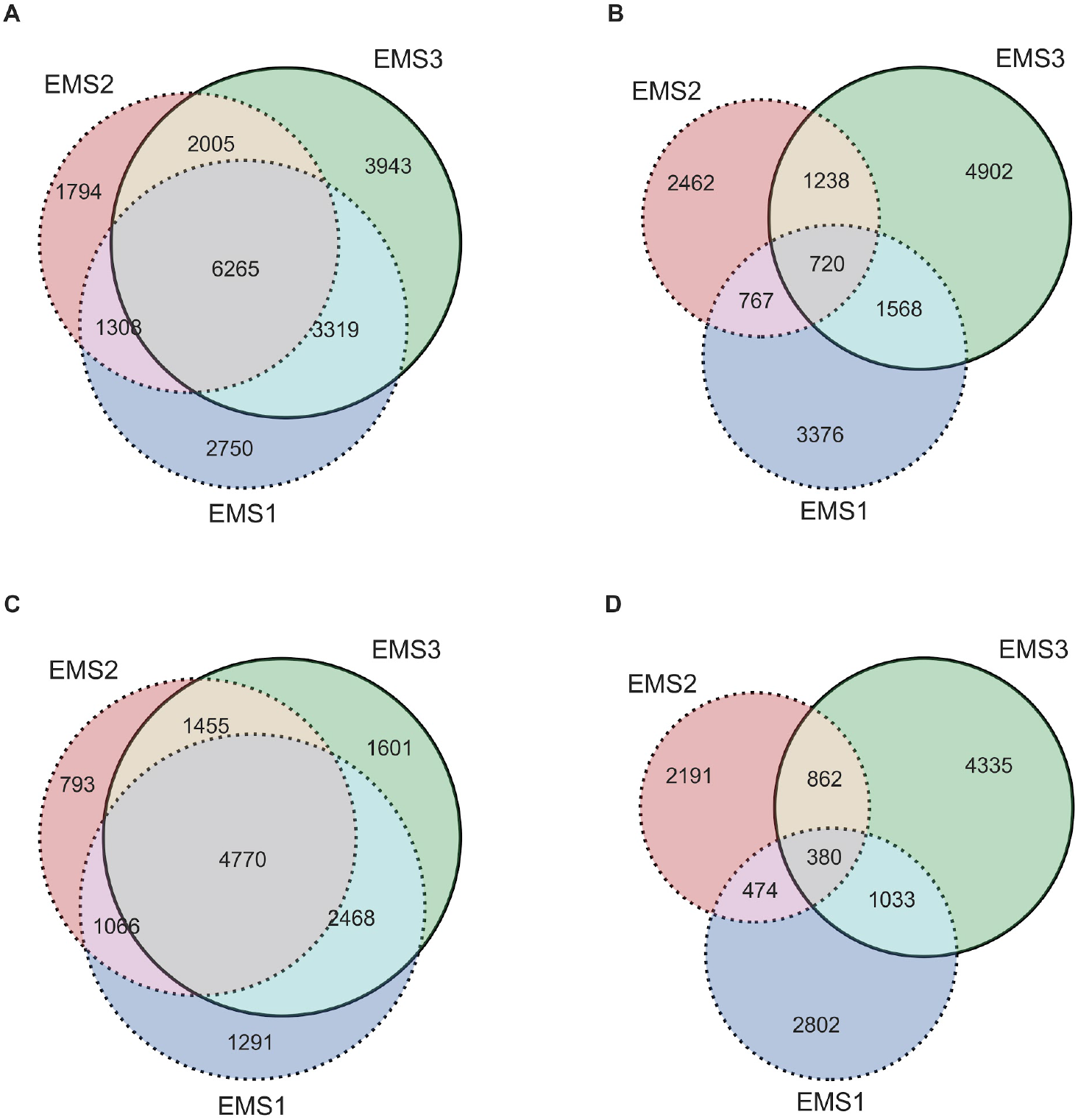
Comparison of the number of significant genes among three datasets. (A) Venn diagrams display the number of shared genes carrying any significant mutations among EMS1 (Purdue), EMS2 (ARS), and EMS3 (CSP, this study). (B) Number of shared genes carrying knockout mutations. (C) Number of shared genes carrying mutations predicted to be significant by SIFT (<0.05) and GERP scores (>=2). (D) Number of shared genes carrying stop-gained mutations.

The distribution of the number of significant mutations in each sample is shown in Figure 2 for EMS datasets and Figure 3A for the natural population. In general, EMS2 and EMS3 carry more significant mutations per sample than EMS1. As discussed above, this might be due to the fact that both EMS2 and EMS3 samples are a mixture of 20 M3 plants while EMS1 is a single M3 plant. The distributions in Figure 2 show that, even for stop-gained mutations, the number of affected genes varies from 0 to 30 per sample for EMS1 (average 10), 4 to 42 for EMS2 (average 20), and 0 to 55 for EMS3 (average 19). These numbers are high but still much smaller than the majority of NP samples with 180 to 420 stop-gained mutations per sample (average 300), as shown in Figure 3A. Less significant genes per sample means less work and less confounding factors in the molecular identification of genes responsible for mutant phenotypes. Therefore, even without considering the large structural variations in the natural population, the BTx623-based EMS data are more efficient resources in elucidating gene functions for forward genetics approaches.

**Figure 2.**
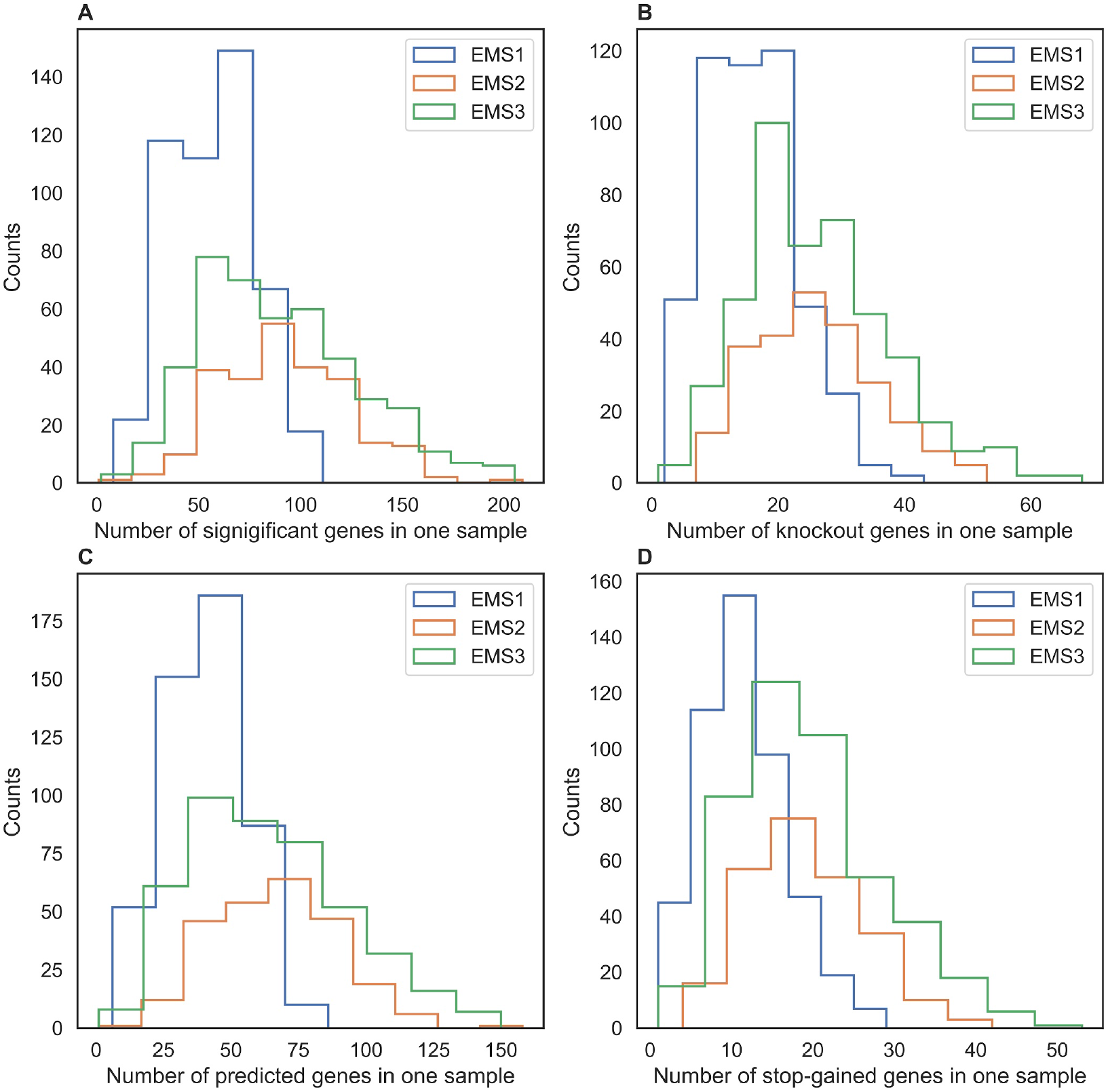
Comparison of the distribution of the number of significant genes in each sample among three EMS datasets. (A) Histograms display the number of genes carrying any significant mutations in each sample for EMS1 (Purdue), EMS2 (ARS), and EMS3 (CSP). (B) Number of genes in each sample carrying blockout mutations. (C) Number of genes in each qsample carrying mutations predicted to be significant by SIFT (<0.05) and evolutionarily conserved by GERP scores (>=2). (D) Number of genes in each sample carrying stop-gained mutations.

**Figure 3.**
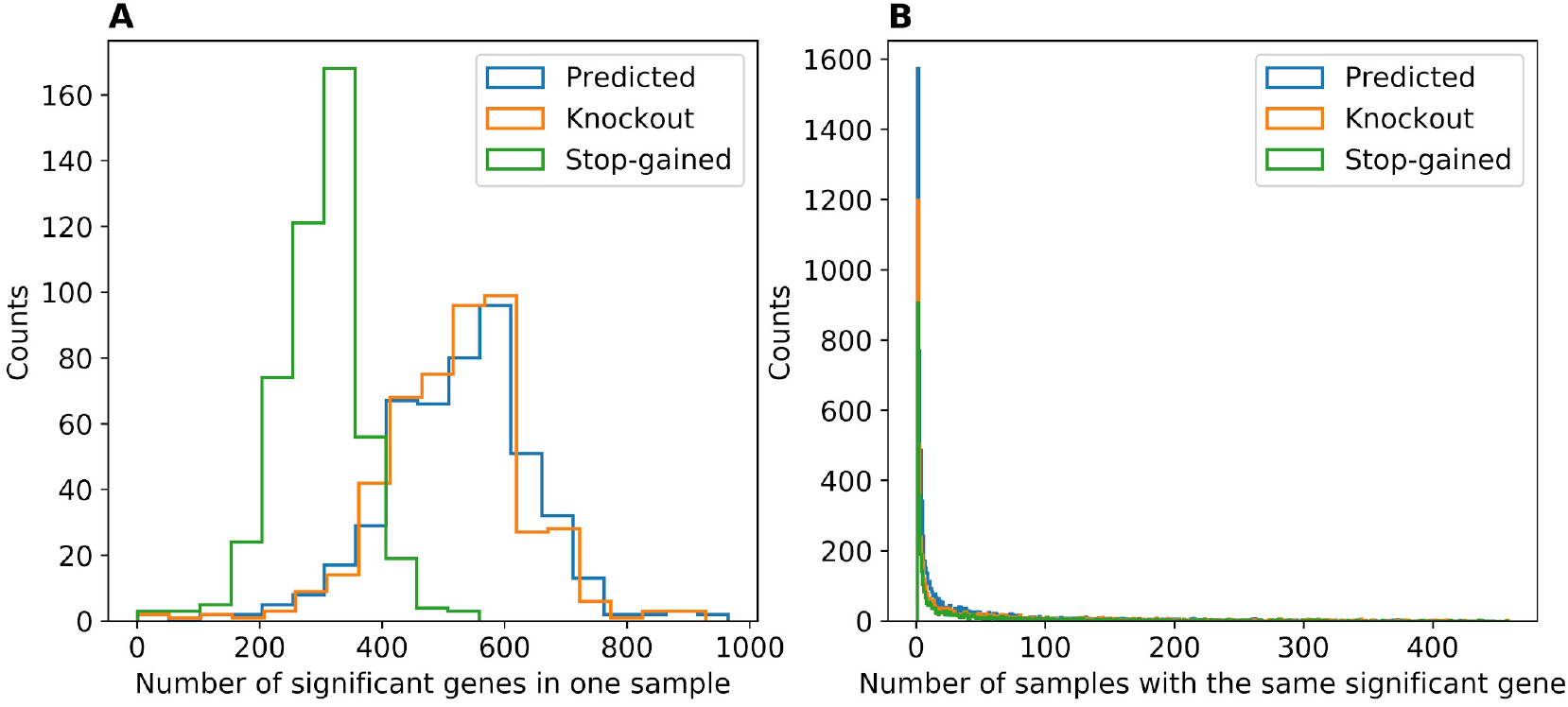
Statistics of significant genes for the natural population. (A) Histograms display the number of genes carrying different kinds of significant mutations in each sample. (B) Number of samples with the same gene carrying different kinds of significant mutations.

As shown in Table 1, there are less genes with stop-gained mutations in the natural population than the EMS populations combined (3550/499 vs 12077/1182). The fact that each sample carries more stop-gained genes than the EMS data is due to the population structure or relatedness among samples. As shown in Figure 3B and Figure 4D, a large number of genes with significant mutations, knockout mutations, or stop-gained mutations, are present in over 25 samples. Some are present in over 250 samples, half of the natural population. On the contrary, the majority of the genes with knockout/stop-gained mutations are present in only one or two EMS samples, with very few present in more than three EMS samples, as shown in Figure 4A, 4B, 4C and Figure S4.

**Figure 4.**
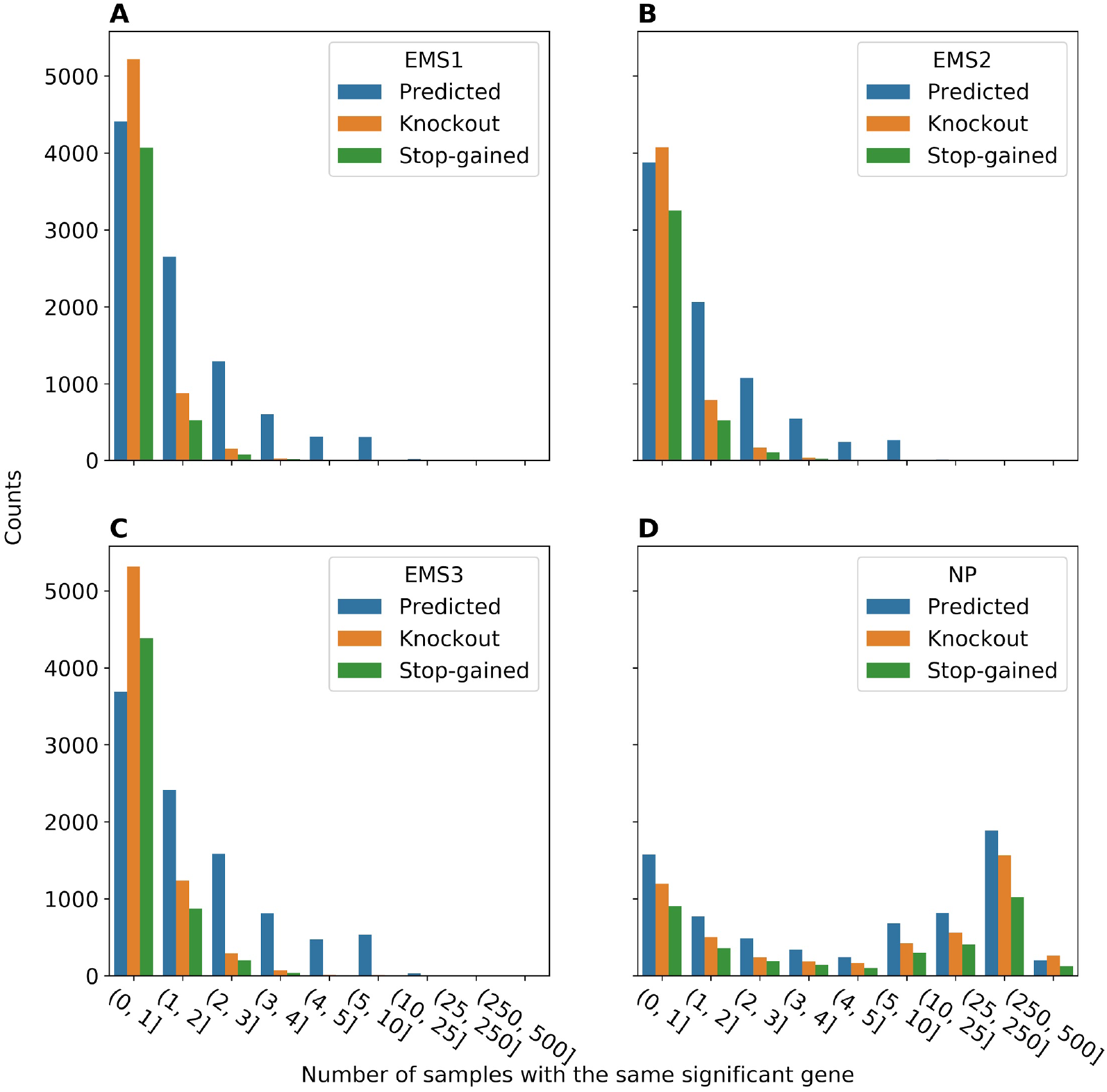
Comparison of distribution of the number of samples with the same significant gene among four datasets. (A) Histograms display the number of samples with the same gene carrying different types of significant mutations for EMS1 (Purdue). (B) Number of samples with the same gene carrying significant mutations for EMS2 (ARS). (C) Number of samples with the same gene carrying significant mutations for EMS3 (CSP). (D) Number of samples with the same gene carrying significant mutations for NP (natural population). A mutation is predicted to be significant by both SIFT (<0.05) and evolutionarily conserved by GERP scores (>=2).

In summary, compared with the natural population, the EMS data carries 4 times more significant mutations (84.8K vs 21.3K), and impacts two times more genes (21.4K vs 11.6K) with any significant mutations, 3 times more genes with knockout mutations (15K vs 5.5K), or four times more genes with stop-gained mutation (12.1K vs 3.6K). On the other hand, each EMS sample carries 6 times less (100 vs 600) genes with any significant mutations, or 20 times less (30 vs 600) genes with knockout mutations, or 15 times less (20 vs 300) genes with stop-gained mutations. Furthermore, the majority of the significant mutations are presented in less than 3 EMS samples in contrast to a large number of them presenting in over 25 samples from the natural population. This reaffirms that the EMS data is more efficient in gene function study.

### Mutation landscape of sorghum

In Figure 5, we combined all three EMS datasets, divided the sorghum genome into non-overlapping 500 kb windows, and calculated the number of significant genes in each window. For comparison, statistics of the significant mutations from the natural population is calculated and shown in blue.

**Figure 5.**
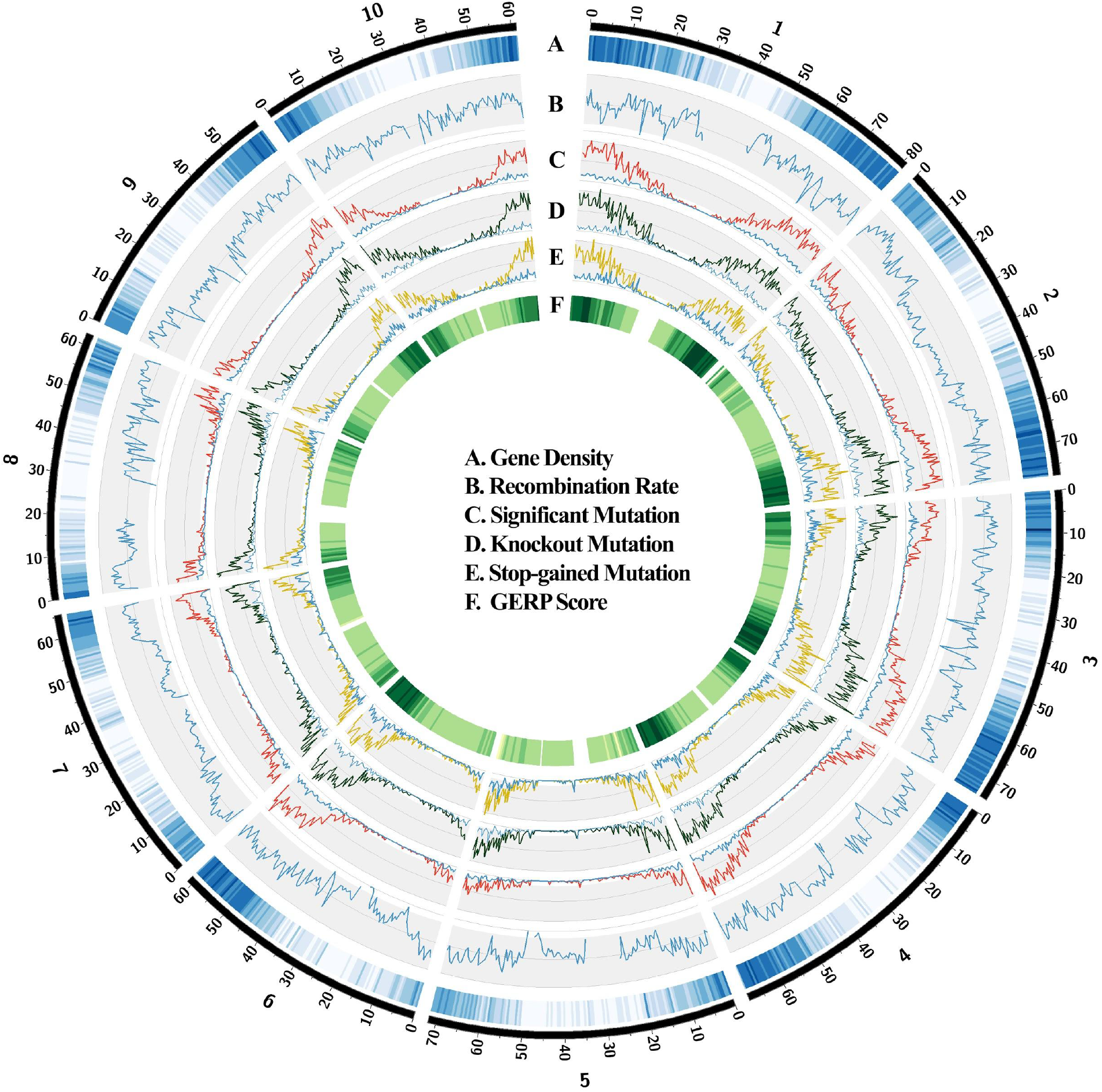
Genome-wide distribution of significant genes in 500 kb windows. (A) Gene density heatmap. (B) Population recombination rates (ρ). (C) Number of genes (minimum 0, maximum 263) with any significant mutations for EMS (red) and natural population (blue). (D) Number of genes (0 - 67) with knockout mutations for EMS (black) and natural population (blue). (E) Number of genes (0 - 51) with stop-gained mutations for EMS (yellow) and natural population (blue). (F) Average GERP scores.

The EMS mutations have been shown to be uniformly distributed within the sorghum genome (Jiao et al. 2016). Figure 5 shows that the number of significant mutations increases with gene densities but is unrelated to the recombination rate. For example, the region around 70 Mb in chromosome 1 has a relatively low recombination rate but no dropping is observed for the number of significant genes from the EMS data. The functions of genes within these regions might be difficult to study with a map-based cloning approach but it is not the case with the EMS method. Inside each 500-kb window, the number of significant, knockout, and stop-gained mutations from the EMS data are much higher than those from the natural population.

## Discussion

With the combined EMS variants (8,409,135 SNPs), there are only 75,457 SNPs present in the natural population (13,087,170 SNPs). This observation suggests that 99.1% of EMS-induced mutations are novel variations that could be used to accelerate sorghum breeding.

Previously, we have established an automated workflow for identification of causal genes through bulk segregation analysis (Wang, Lu, Regulski, et al. 2020). We built a database with the EMS2 data to support complementary testing of candidate causal genes. EMS lines with alternative significant mutations on the same gene can be retrieved from the database and used to verify the true causal relationship between the gene and the observed mutant phenotype. In this study, we have integrated all three sorghum EMS datasets into the database, which increases the gene coverage from 33.3% (11,372) to 62.7% (21384), as shown in Table 1. If using SIFT score < 0.05 as the only criteria for a mutation to be predicted as significant (Jiao et al. 2016) (without considering median_info and GERP score), the total number of genes with significant mutations increases from 20561 (or 60.3%) to 29478 (or 86.4%) after combining the EMS1 and EMS3 data. Considering that loss of some genes might be lethal to the plant, we have closely covered the majority of the sorghum gene space. By building a massive scale of genomic resources for the sorghum community, we hope to facilitate identifying and validating gene functions with mutant phenotypes within a short period.

---

**Figure S1.**
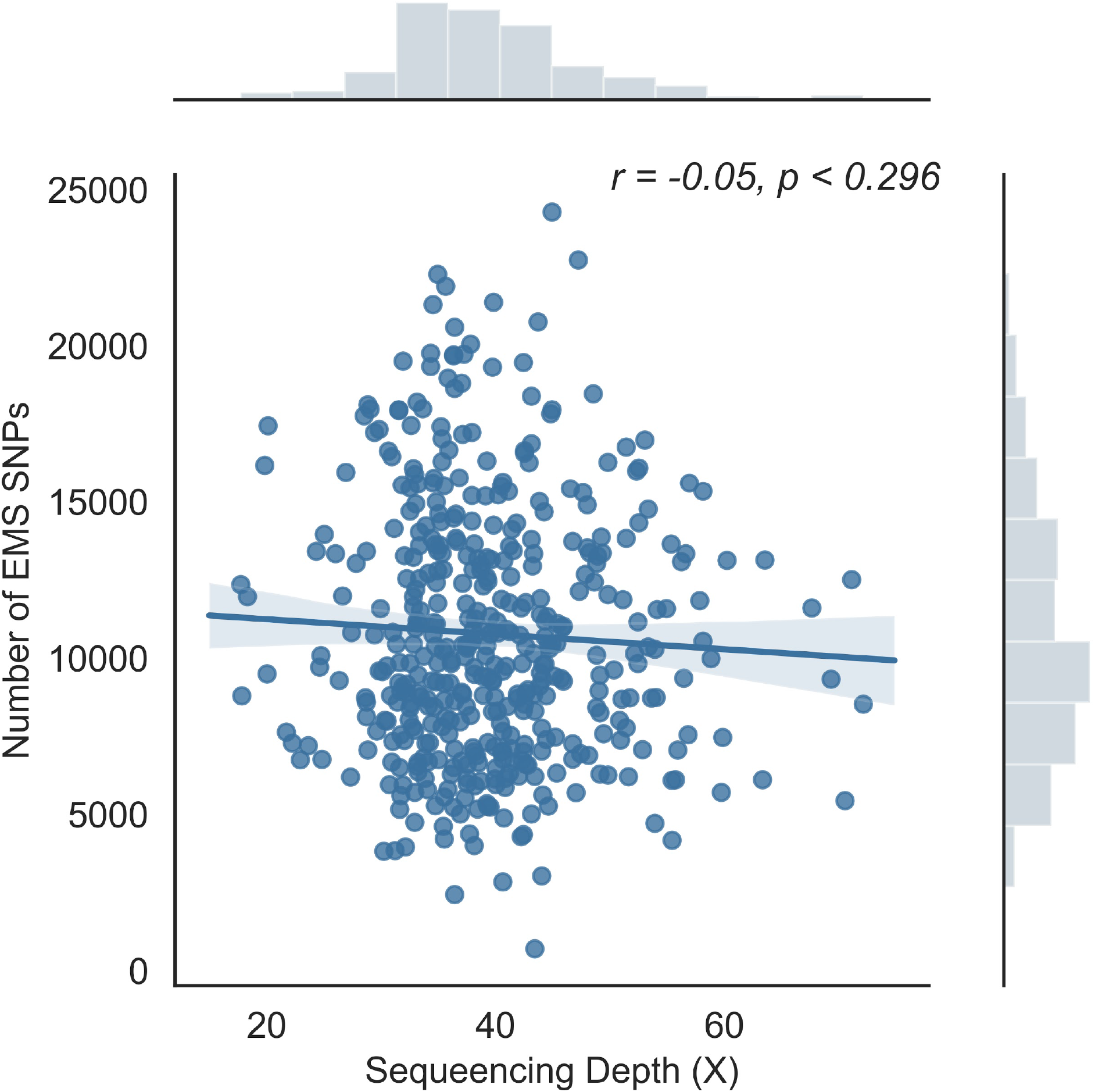
Relationship between the number of EMS mutations with sequencing depth for the EMS3 data set. The outlier with significantly more EMS SNPs (ARS1162, with 54085 EMS SNPs) is removed.

**Figure S2.**
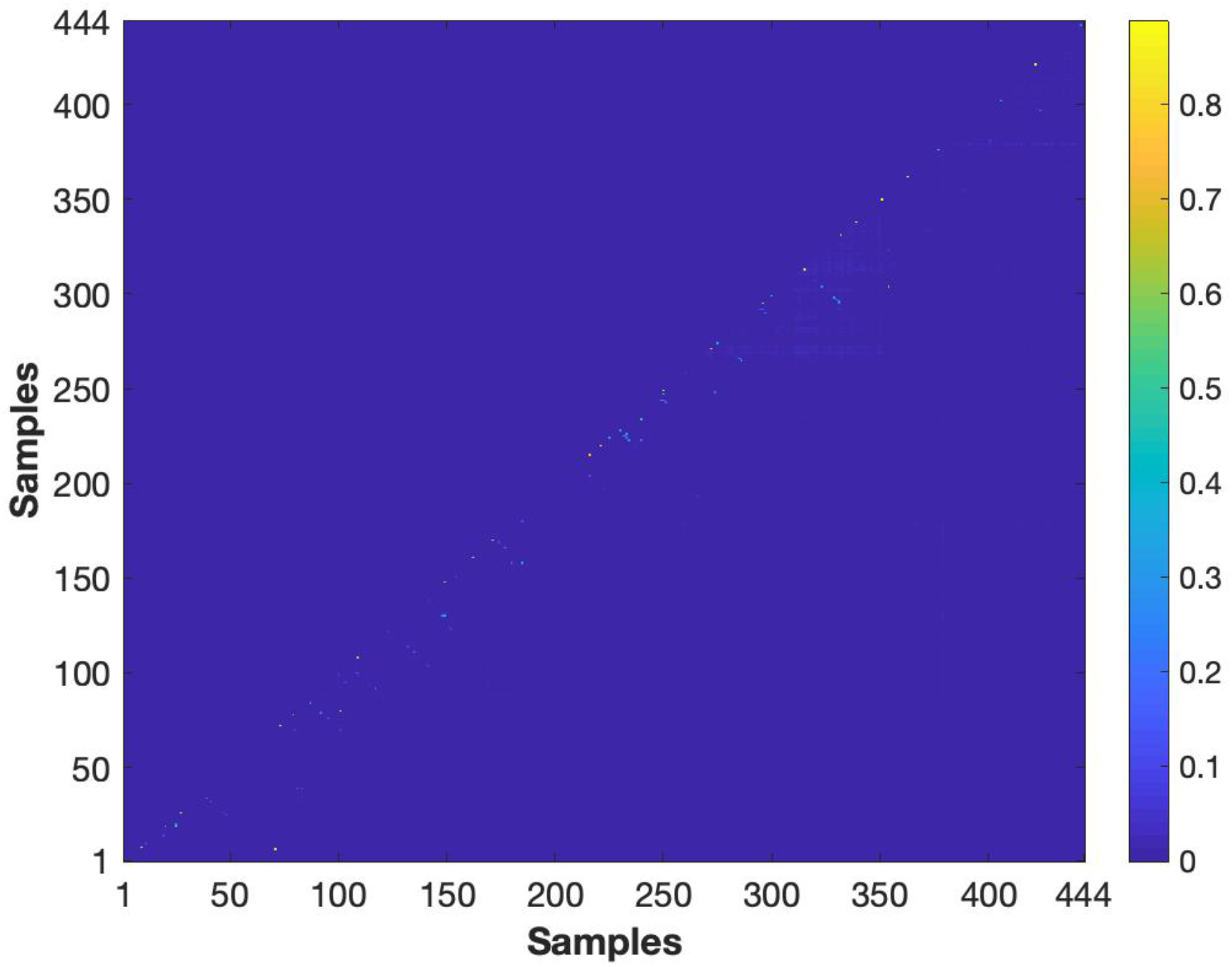
Percentage of the overlapping EMS SNPs between each pair of samples for the EMS3 data set. 99.9% of the pairwise comparisons have < 5% identical EMS mutations, in which 97% have <1% identical EMS mutations.

**Figure S3.**
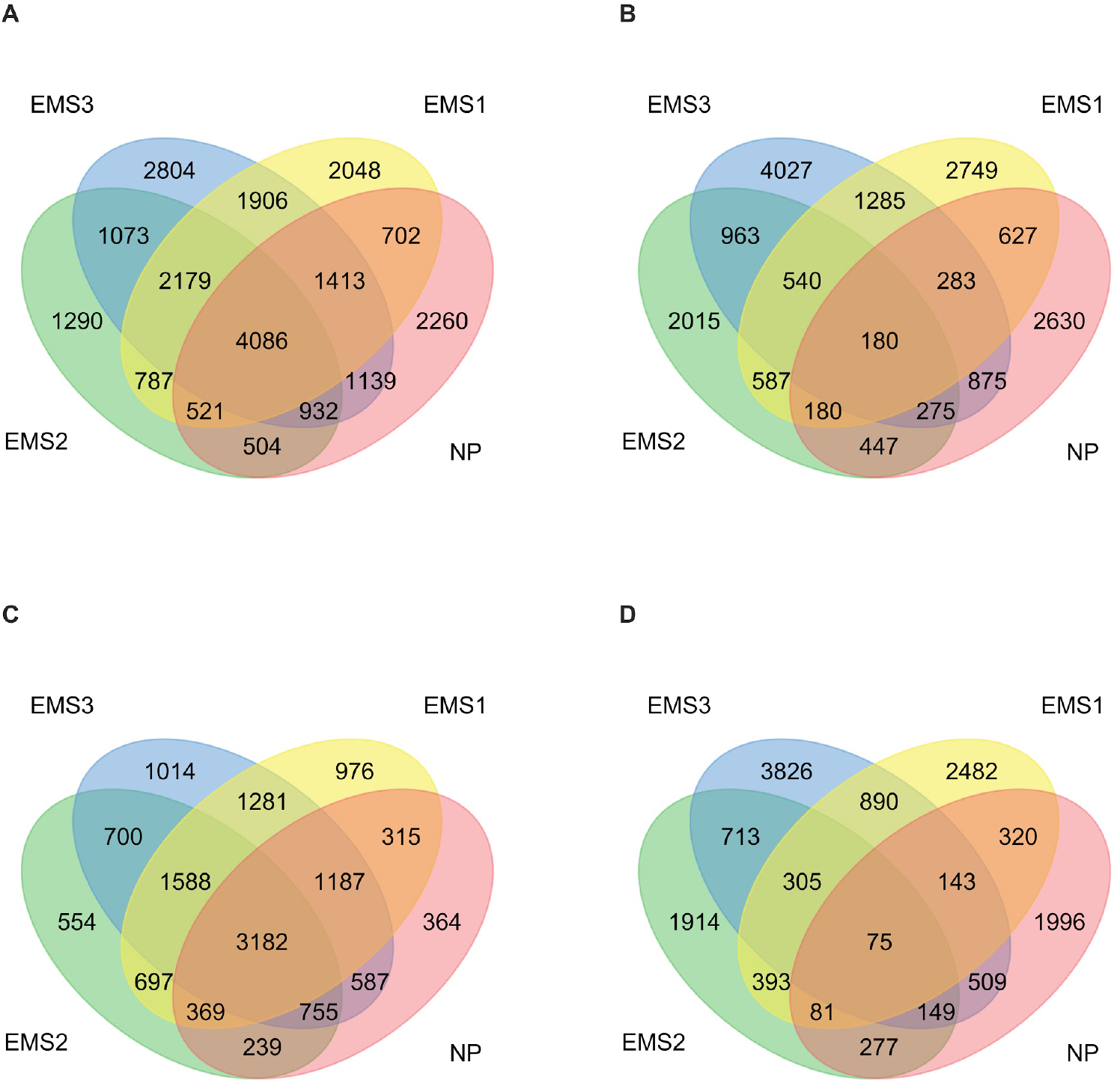
Comparison of the number of significant genes among four datasets. (A) Venn diagrams display the number of shared genes carrying any significant mutations among EMS1 (Purdue), EMS2 (ARS), EMS3 (CSP), and the NP (the natural population). (B) Number of shared genes carrying blockout mutations. (C) Number of shared genes carrying mutations predicted to be significant by SIFT (<0.05) and evolutionarily conserved by GERP scores (>=2). (D) Number of shared genes carrying stop-gained mutations.

**Figure S4.**
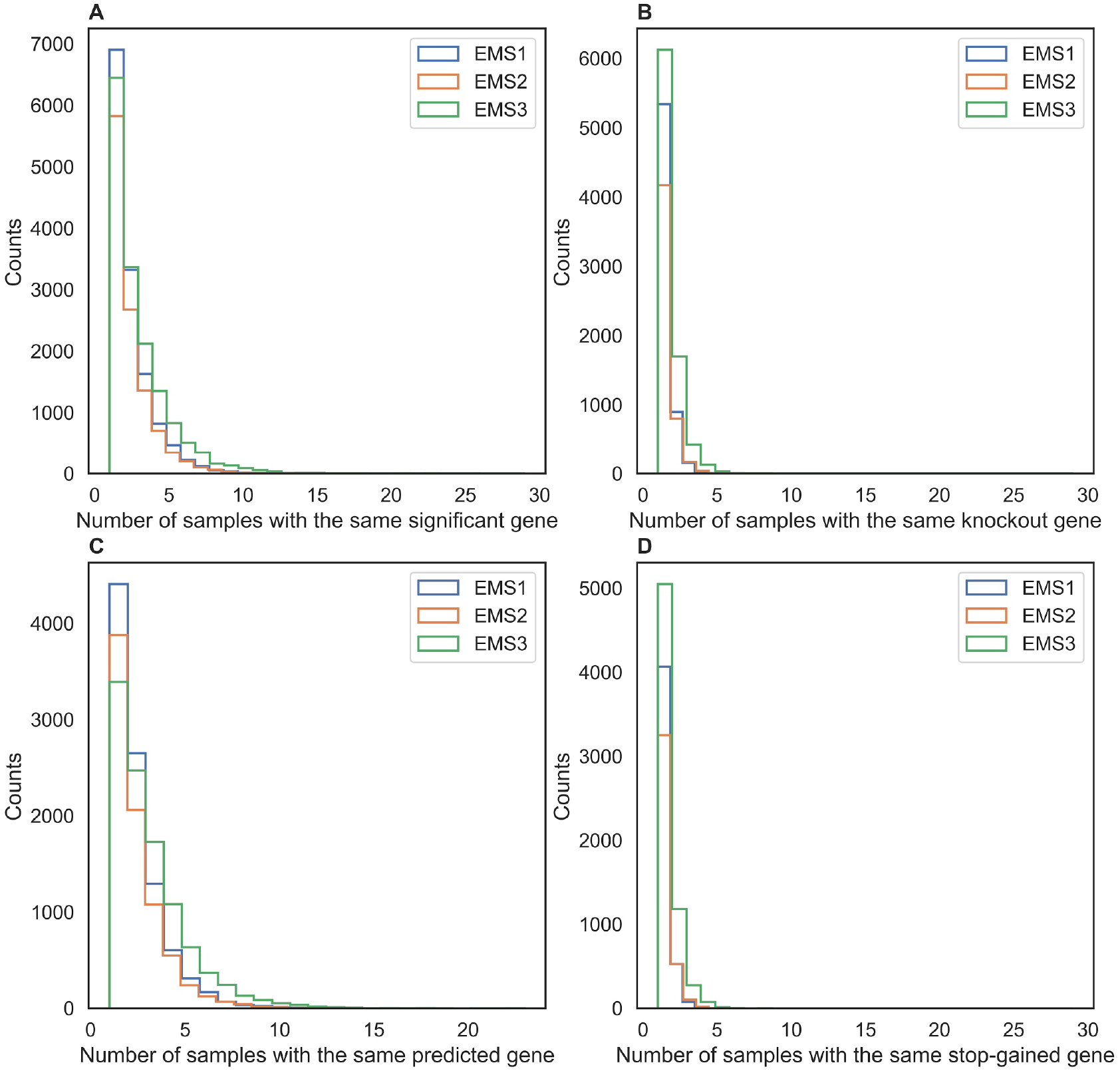
Comparison of distribution of the number of samples with the same significant gene among three datasets. (A) Histograms display the number of samples with the same gene carrying any significant mutations for EMS1 (Purdue), EMS2 (ARS), and EMS3 (CSP). (B) Number of samples with the same gene carrying blockout mutations. (C) Number of samples with the same gene carrying mutations predicted to be significant by SIFT (<0.05) and evolutionarily conserved by GERP scores (>=2). (D) Number of samples with the same gene carrying stop-gained mutations.

## References

Abe, Akira, Shunichi Kosugi, Kentaro Yoshida, Satoshi Natsume, Hiroki Takagi, Hiroyuki Kanzaki, Hideo Matsumura, et al. 2012. “Genome Sequencing Reveals Agronomically Important Loci in Rice Using MutMap.” Nature Biotechnology 30 (2): 174–78.

Addo-Quaye, Charles, Elizabeth Buescher, Norman Best, Vijay Chaikam, Ivan Baxter, and Brian P. Dilkes. 2017. “Forward Genetics by Sequencing EMS Variation-Induced Inbred Lines.” G3 7 (2): 413–25.

Addo-Quaye, Charles, Mitch Tuinstra, Nicola Carraro, Clifford Weil, and Brian P. Dilkes. 2018. “Whole-Genome Sequence Accuracy Is Improved by Replication in a Population of Mutagenized Sorghum.” G3 8 (3): 1079–94.

Cingolani, Pablo, Adrian Platts, Le Lily Wang, Melissa Coon, Tung Nguyen, Luan Wang, Susan J. Land, Xiangyi Lu, and Douglas M. Ruden. 2012. “A Program for Annotating and Predicting the Effects of Single Nucleotide Polymorphisms, SnpEff: SNPs in the Genome of Drosophila Melanogaster Strain w1118; Iso-2; Iso-3.” Fly 6 (2): 80–92.

Davydov, Eugene V., David L. Goode, Marina Sirota, Gregory M. Cooper, Arend Sidow, and Serafim Batzoglou. 2010. “Identifying a High Fraction of the Human Genome to Be under Selective Constraint Using GERP++.” PLoS Computational Biology 6 (12): e1001025.

Jiao, Yinping, John Burke, Ratan Chopra, Gloria Burow, Junping Chen, Bo Wang, Chad Hayes, Yves Emendack, Doreen Ware, and Zhanguo Xin. 2016. “A Sorghum Mutant Resource as an Efficient Platform for Gene Discovery in Grasses.” The Plant Cell 28 (7): 1551–62.

Li, Heng, and Richard Durbin. 2009. “Fast and Accurate Short Read Alignment with Burrows-Wheeler Transform.” Bioinformatics 25 (14): 1754–60.

Li, Heng, Bob Handsaker, Alec Wysoker, Tim Fennell, Jue Ruan, Nils Homer, Gabor Marth, Goncalo Abecasis, Richard Durbin, and 1000 Genome Project Data Processing Subgroup. 2009. “The Sequence Alignment/Map Format and SAMtools.” Bioinformatics 25 (16): 2078–79.

Lozano, Roberto, Elodie Gazave, Jhonathan P. R. Dos Santos, Markus G. Stetter, Ravi Valluru, Nonoy Bandillo, Samuel B. Fernandes, et al. 2021. “Comparative Evolutionary Genetics of Deleterious Load in Sorghum and Maize.” Nature Plants 7 (1): 17–24.

McCormick, Ryan F., Sandra K. Truong, Avinash Sreedasyam, Jerry Jenkins, Shengqiang Shu, David Sims, Megan Kennedy, et al. 2018. “The Sorghum Bicolor Reference Genome: Improved Assembly, Gene Annotations, a Transcriptome Atlas, and Signatures of Genome Organization.” The Plant Journal: For Cell and Molecular Biology 93 (2): 338–54.

Page, Damian R., and Ueli Grossniklaus. 2002. “The Art and Design of Genetic Screens: Arabidopsis Thaliana.” Nature Reviews. Genetics 3 (2): 124–36.

Valluru, Ravi, Elodie E. Gazave, Samuel B. Fernandes, John N. Ferguson, Roberto Lozano, Pradeep Hirannaiah, Tao Zuo, et al. 2019. “Deleterious Mutation Burden and Its Association with Complex Traits in Sorghum (Sorghum Bicolor).” Genetics 211 (3): 1075–87.

Vaser, Robert, Swarnaseetha Adusumalli, Sim Ngak Leng, Mile Sikic, and Pauline C. Ng. 2016. “SIFT Missense Predictions for Genomes.” Nature Protocols 11 (1): 1–9.

Wang, Liya, Zhenyuan Lu, Melissa delaBastide, Peter Van Buren, Xiaofei Wang, Cornel Ghiban, Michael Regulski, et al. 2020. “Management, Analyses, and Distribution of the MaizeCODE Data on the Cloud.” Frontiers in Plant Science 11 (March): 289.

Wang, Liya, Zhenyuan Lu, Michael Regulski, Yinping Jiao, Junping Chen, Doreen Ware, and Zhanguo Xin. 2020. “BSAseq: An Interactive and Integrated Web-Based Workflow for Identification of Causal Mutations in Bulked F2 Populations.” Bioinformatics, August. https://doi.org/10.1093/bioinformatics/btaa709.

Wang, Liya, Zhenyuan Lu, Peter Van Buren, and Doreen Ware. 2018. “SciApps: A Cloud-Based Platform for Reproducible Bioinformatics Workflows.” Bioinformatics 34 (22): 3917–20.

Xin, Zhanguo, Ming Li Wang, Noelle A. Barkley, Gloria Burow, Cleve Franks, Gary Pederson, and John Burke. 2008. “Applying Genotyping (TILLING) and Phenotyping Analyses to Elucidate Gene Function in a Chemically Induced Sorghum Mutant Population.” BMC Plant Biology 8 (October): 103.

